# Transcutaneous delivery of disease-specific PI3K/Akt/mTOR inhibitor-based hybrid nanoparticles in hydrogel system for the management of psoriasis: Insights from *in vivo* studies

**DOI:** 10.1101/2024.06.17.599287

**Authors:** Roshan Keshari, Rupali Bagale, Sulagna Rath, Abhijit De, Rinti Banerjee, Shamik Sen, Rohit Srivastava

## Abstract

We have reported robust and scalable lipid polymeric conjugated hybrid nanoparticles comprised of phospholipid shell and polymeric core that combine the benefit of both polymeric and lipid nanoparticles. Rapamycin, a PI3K/Akt/mTORC1 inhibitor, was encapsulated inside the lipid-polymeric conjugated spherical-shaped hybrid nanoparticles (RPMN), having an encapsulation efficiency of ≈ 83% and particle size ≈ 277.6 nm. Further, RPMN was converted into the carpool-based hydrogel system (RPMNGel), which enhanced release kinetics, long-term stability and skin residence time. Specifically, in an *in-vivo* imiquimod-induced psoriatic model, RPMNGel showed high accumulation and deeper penetration inside the epidermis, slowly diffusing away inside the psoriatic skin without causing any side effect to normal skin. This leads to longer and sustained retention over more than three days without being affected by sweat, humidity or wiping due to adherence between the stratum corneum and epidermis. Similarly, the cumulative PASI score was also reduced from 10.25 to 1.75 on day 7 in the group treated with RPMNGel. Overall, RPMNGel has a potential role in treating and managing psoriasis.

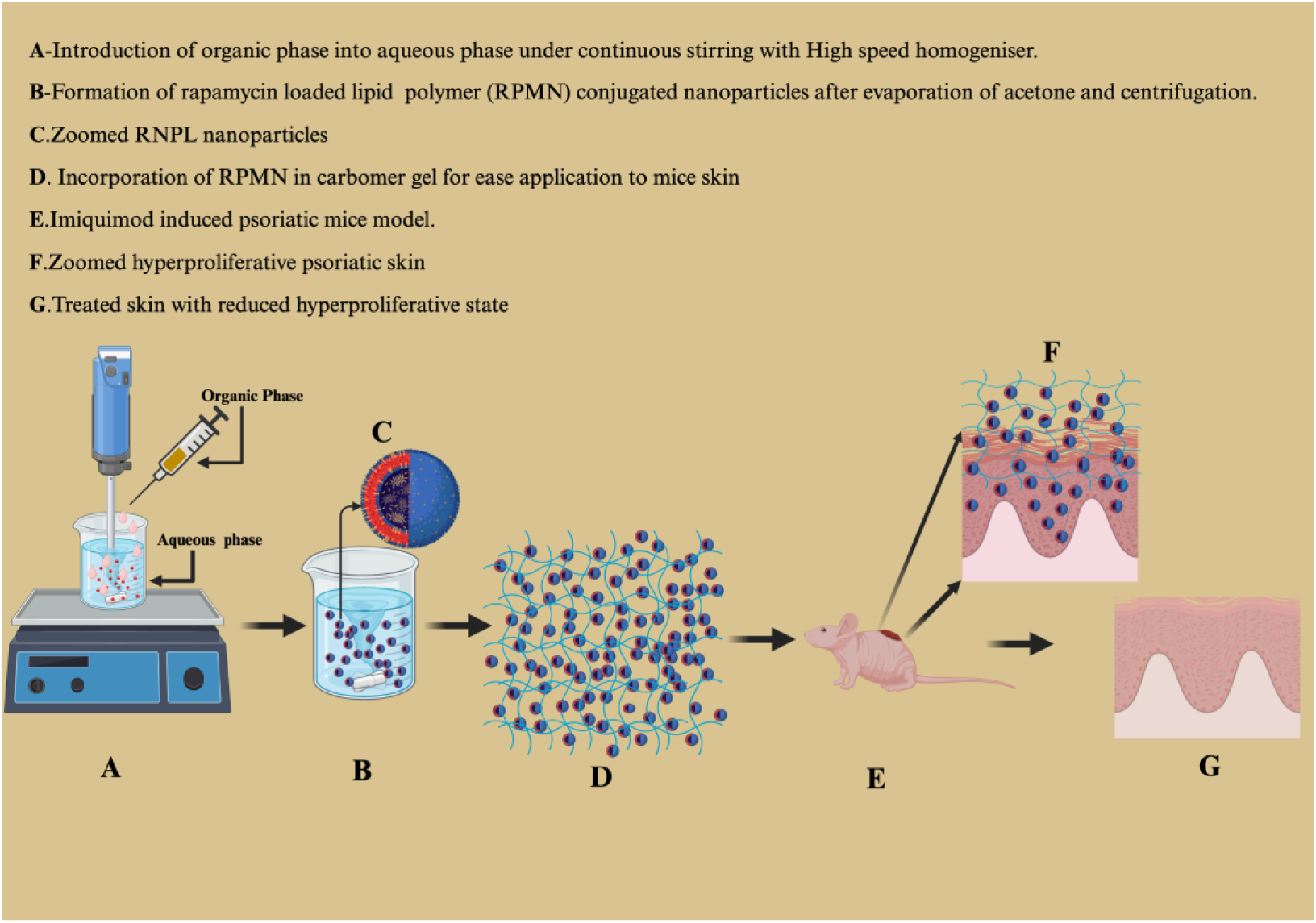

## 1. Introduction

Psoriasis is a recurring immune skin disorder characterised by epidermal acanthosis, over-proliferation of the keratinocytes, alteration of epidermal differentiation, dermal angiogenesis and presence of dense inflammatory cells [1]. Clinical symptoms include scaly, itchy, painful plaque, cracking, bleeding, and distorted skin [2]. It affects around 2% of the global population and negatively impacts the quality of life[3]. The exact cause and mechanism are still unclear. Still, it is assumed that it might be due to stimulating immune cells, such as T cells, neutrophils, dendritic cells, and macrophages, which release substances that cause hyperproliferation, thickening, and angiogenesis [4]. While many hypotheses exist, the most significant evidence is the association of genes like PSORS1, IL-23R, and IL-12R[5]. Despite the availability of different treatment regimens like systemic, topical, and phototherapy, none of them is competent to cure the disease, and they can only reduce the symptoms to a large extent, which poses a significant challenge to developing more effective and better ways of treatment.

Recently, attention has been paid to the PI3K/Akt/mTORC1 signalling cascade as a crucial homeostatic epidermal regulator that may be involved in inflammatory skin conditions [6]. Previously, it was shown that psoriatic lesions exhibit hyper-activation of mTOR kinase and its downstream target, ribosomal protein S6 [7,8] This pathway may also contribute to the imbalance of Th1, Th2 and Th17 cells in psoriasis [9]. Similarly, Akt activation has also been observed in psoriatic lesions, including the basal Ki-67+ proliferating cells and all epidermal layers [10,11]. This might be because psoriatic keratinocytes continue proliferating after leaving the basal layer.

Consequently, hyperproliferation of epidermal keratinocytes promotes psoriasis by activating the PI3K/Akt/mTOR signalling pathway. Rapamycin (PI3K/Akt/mTOR signalling pathway inhibitor), also known as sirolimus, works when cytoplasmic protein FKPB-12, binds to rapamycin, interacts with mTORC1 to inhibit the mTOR target proteins S6 and 4E-BP, which are necessary for protein synthesis and, consequently, for the growth and proliferation of cells [12]. Thus, we used imiquimod to induce psoriatic-like skin lesions to investigate the therapeutic efficacy of rapamycin and understand its role in managing psoriasis. Histologically, IMQ treatment causes keratinocyte hyperproliferation, aberrant epidermis differentiation, elevated angiogenesis, and a high level of immune cell infiltration [13,14]. Application of imiquimod in mice model has shown the activation and upregulation of the PI3K-mTOR cascade which are comparable to psoriasis in humans [15] and can be relieved by inhibiting this pathway [16]. These results highlight the importance of the mTOR pathway in the pathophysiology of psoriasis and strongly support topical rapamycin as an anti-psoriatic agent.

Here, we have prepared lipid polymeric conjugated hybrid nanoparticles (LPCHN) comprised of a phospholipid shell and polymeric core that combine the benefit of both polymeric and liposomal nanoparticles [17,18]. Both have their advantages; monolayer lipids provide stability by restricting the outward diffusion of the encapsulated drugs, and polymeric cores enhance the encapsulation of poorly water-soluble drugs like rapamycin [19]. Several lipid, polymer and their ratio were evaluated to develop a stable LPCHN that can penetrate the deeper layer of the skin with a controlled release rate.

This study is designed to measure the usefulness of topical application rapamycin-loaded LPCHN known as RPMN in an imiquimod-induced psoriasis mice, focusing on reducing disease progression, permeation and deposition of a drug at various layers.

## 2. Materials and Method

Poly(lactic-co-glycolic acid) copolymer of DL-lactide and Glycolide in a 50:50 molar ratio (MW-7,000 – 17,000 da) were purchased from Nomisma Healthcare India; Phospholipon 90G (p@90G) was obtained from Lipoid GmbH Germany; Rapamycin (99.4% purity) were purchased from MedChem Express; polyvinyl alcohol (mw approx.1,15000) were procured from Loba Chemie, India; dialysis membrane (12.5 KDa), fetal bovine serum (FBS), antibiotic-antimycotic solution, 3-(4,5-dimethylthiazol-2-yl) −2,5-diphenyl tetrazolium bromide (MTT) reagent, Dulbecco’s modified eagle medium (DMEM), dimethyl sulfoxide (DMSO) and phosphate buffer saline (PBS) pH 7. 4 were purchased from HImedia laboratories; Tocopherol polyethylene glycol 1000 succinate (TPGS) was procured from matrix science private limited; tween-80, triethanolamine, and carbopol 974P were purchased from Sigma Aldrich. AddexBio supplied HaCaT cells; Rhodamine 6G (≥97%) were purchased from Merck KGaA. Imiquimod was purchased (5% W/W; Glenmark Pharmaceuticals, India) from a local market. In every experiment, high-purity deionized water (resistivity 18.2 M cm) purified by a Milli-Q Plus water purification system was used (Millipore, USA). HaCaT cells were grown as an adherent monolayer at 37°C in 5% CO2 in DMEM enriched with 10% FBS and 1% antibiotic. After attaining a confluency of 70–80%, cells were subcultured.

### 2.1 Formulation of rapamycin-loaded lipid polymeric conjugated hybrid nanoparticles (RPMN)

The lipid-polymeric conjugated hybrid nanoparticles were prepared using the emulsification and solvent evaporation method described by Gurny et al. [20]. Briefly, a solution of PVA as an aqueous phase (2mg/ml) was prepared previously, and P@90G (5 mg/ml) was added to the PVA solution and transferred to a magnetic stirred at 500-600 RPM at 37°C and when it solubilized TPGS (5mg/ml) and tween 80 (2mg/ml) were added and allowed to mix until it gets completely solubilize. Separately, PLGA (15mg/ml) were dissolved in acetone on a magnetic stirrer at RPM of 600-800 until completely dissolved. When PLGA was dissolved entirely, rapamycin (6mg/ml) was accurately weighed and added to the PLGA solution. The prepared mixture was continuously stirred for another 3 hours at 400-500 RPM. When both the aqueous and organic phases were ready, the organic phase solution was introduced into the aqueous phase drop by drop slowly with continuous stirring at RPM of 4000 by using a high-speed homogenizer. After that, the formulation was transferred to a magnetic stirrer at RPM of 1200-1400 RPM at room temperature till the evaporation of organic solvent. The final lipid polymeric conjugated nanoparticles were centrifuged at 10,000 RPM for 15 min at 4 °C; the supernatant was discarded, washed twice with Milli-Q water, and suspended in Milli-Q water.

### 2.2. Incorporation of lipid-polymeric conjugated nanoparticles to the hydrogel system (RPMNGel)

Prepared RPMN were further incorporated into (0.8% w/v) Carbopol 974P to prepare a hydrogel system. Briefly, Carbopol 974 P were dispersed in RPMN for 12 hours at continuous stirring (1000-1200 RPM) to form a like gel. Further, it was again stirred vigorously by a high-speed homogenizer to form a uniform mixture. The gel was made alkaline (pH 6.5) using triethanolamine (6.5 µl/ ml). A texture analyser and rheometer further analysed the texture and viscosity of the prepared gel.

### 2.3 Particle size distribution, TEM Imaging, Drug entrapment efficiency

The dynamic light scattering (DLS) method was used (Malvern Zeta sizer Malvern Instrument Ltd., UK) to determine the Z-average, polydispersity index (PDI), and zeta potential of RPMN. The linked functional data from laser scattering were carefully fitted, and the Z-average size was calculated using the average size as an intensity value. The formulation was diluted 100 times in double-distilled water before analysis.

RPMNs’ morphological characteristics were captured using transmission electron microscopy (TEM). The copper grid (300 mesh) was coated with a drop of diluted RPMN and dried at room temperature. A drop of phosphotungstic acid (2%) was applied and dried for 30 minutes before being scanned using a 300 V TEM (FEI Tecnai G2, F30).

The entrapment efficiency of RPMN was studied using an ultra-centrifugation technique. RPMN was centrifuged (Sigma 4K15, Refrigerated high-speed benchtop centrifuge) at 10,000 RPM for 15 minutes at four °C, and the supernatant was discarded. The resulting pellet was washed twice with Milli-Q water before being reconstituted in Milli-Q water. The reconstituted formulation was further diluted ten times with methanol, and drug entrapment efficiency was calculated using HPLC under the chromatographic conditions outlined in point 4. The EE has been calculated as follows:

> **EE-(Drug in nanoparticles/Initial drug added)*100**

### 2.4. *In vitro* drug release

The release kinetics of rapamycin from free rapamycin gel (Free RAGel) and rapamycin-loaded nanoparticle gel (RPMNGel) were investigated utilizing the dialysis bag method with 40% methanol in PBS as a sink medium and a 12.5 kDa dialysis membrane. The desired sample volume was collected at various time intervals, i.e. 0.5, 1, 2, 4, 6, 8, 12, 24, and 48 hours, and the sink medium was substituted with a corresponding volume. The total percentage of rapamycin present was calculated using the HPLC method using the chromatographic condition outlined in point 4.

### 2.5 Stability Studies

Stability studies of RPMNGel were conducted for 1, 2, and 4 months at various storage conditions such as 4°C, ambient temperature, and accelerated environments, especially 40°C and 75% RH. Samples were collected at the above-suggested intervals, and visual observations such as syneresis, uniformity, variation in colour, separation of phases, roughness, and drug content analysis were examined by reconstituting 50mg of RPMNGel in 950µL of methanol and centrifuged at 10,000 RPM for 15 minutes, and the supernatant was analysed via HPLC.

### 2.6 Rheological Behaviour

A rheometer was used to study the rheological characteristics of RPMNGel. A 25 mm diameter parallel disc was used, with the probe and disc kept at a 0.5 mm distance. The shaft was rotated for three minutes at a shear rate of 10 s^−1^ and an equilibration duration of 9 seconds at 25°C. Equivalent viscosities (cP) were measured at 20 time points, and the final viscosity (cP) was calculated using the average of three readings. Continuous shear stress was used to calculate the shear rate (1/s) as a function of shear stress (mPa), with the associated shear stress measured at 37°C.

### 2.8 Texture analysis

The texture analysis of RPMNGel was examined using the TA2/1000 probe on the CT3-1000 Texture Analyzer (Brookfield Engineering Labs, Inc.) in compression mode. The detailed parameter used is mentioned in **(Table Supp. S1)**. Two cone-shaped probes, male and female, were used. The female probe was connected to the sample holder’s base, while the male probe was connected to the load. The gel deformed at a rate of 2.0mm/s with a zero trigger load. The male probe was pulled up at the same speed, and the result was recorded on TexturePro software [21,22].

### 2.9 In *vitro* antioxidant assay

The anti-oxidative and free radical scavenging characteristics of the RPMN have been examined utilizing the DPPH Assay method. A 0.1 mM concentration of DPPH solution was prepared in ethanol. A mixture of 3.9 ml of DPPH and 100 µL RPMN were prepared and incubated for 10 min, and absorbance was recorded at 517 nm[23]. Free radicals antioxidant activities were analyzed using the following formula:

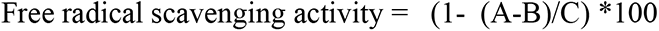

Where,

A: absorbance of DPPH and RNPL

B: Absorbance of RNNPL and

C: Absorbance of DPPH

### 2.10. *In vivo* and *In vitro* cytotoxicity of nanoparticles

The cytotoxicity of RPMN, BNL (Blank nanoparticles), and free rapamycin in DMSO was assessed using an MTT assay. HaCaT cells were seeded at an average density of 10,000 cells/well in a 96-well plate and incubated for 24 hours at 37°C in 5% CO_2_. After 24 hours, cells were rinsed and treated with the above formulations at doses ranging from 5 µg to 200 µg/mL for 24 hours. After treatment, the culture medium was discarded, rinsed with PBS, and MTT (0.5 mg/mL) was added and incubated for 4 hours. After 4 hours, MTT solution was removed, the formazan crystals were dissolved in DMSO, and absorbances were measured at 570 nm with a microplate reader (Tecan Switzerland).

To investigate the in *vivo* toxicity of nanoparticles, 100 μl of either PBS and equivalent to 100 μl RPMNGel, BNLGel or Free RAGel were applied daily to the dorsal mice skin. After 72 hours, the mice were euthanized, dorsal skin was collected, fixed with 10% neutral buffered formalin for at least 24 hours, tissues were embedded in paraffin and sliced into five μm-thick sections, followed by haematoxylin & eosin staining. The sections were observed under Zeiss Axio Imager Z1, Carl Zeiss, Germany. For cytotoxicity assessment, five fields from each representative slice of each mouse were chosen at random.

### 2.11. Ex *vivo* skin penetration and deposition

To study the penetration profile of RPMNGel and free rapamycin gel (Free RAGel) through the skin, extracted skin from psoriatic Swiss albino mice was used [24]. The skin tissue was separated from adipose tissue and attached to a Franz diffusion apparatus between receiver and donor compartment and tightened with a clamp, which had been previously filled with 40% methanol in PBS [25]. RPMNGel and Free RAGel were added to the donor compartment and allowed to penetrate for 24 hours at 37°C under stirring at 500-700 RPM.

After 24 hours, the skin sample was collected, washed thrice with PBS to remove unabsorbed drugs, and dried. The stratum corneum layer was separated using the cellophane tape stripping method. The first two pieces of stripped tape were discarded as they contained unabsorbed drugs. Ten strips were detached and soaked in methanol for 24 hours at 4°C for rapamycin extraction. The remaining skin was cut into small pieces and processed using a tissue homogenizer [26]. The sample was transferred to a shaker at 4°C for 24 hours and then centrifuged and analysed using HPLC, as mentioned in point 4.

## 3. In *vivo* study on imiquimod induced psoriatic mice model

### 3.1 In *vivo* transcutaneous delivery of nanoparticles inside psoriatic skin

The in *vivo* penetration of disease-free+ Free R6G@Gel, disease–free+ R6G@RPMNGel, psoriasis+ Free R6G@Gel, psoriasis+ R6G@RPMNGel were visualized by using rhodamine 6G (R6G) fluorescent dye on the ear of IMQ exposed mice. Briefly, R6G@ RPMNGel was prepared same as RPMNGel. The ear of IMQ-exposed mice was washed, gently wiped, dried, and then above-mentioned dye-loaded formulation was gently applied on the affected ear skin and allowed to penetrate for 24 hours. After 24 hours, mice were euthanized, ear skin was collected, unabsorbed formulations were removed by multiple washing with PBS and visualized under a laser scanning confocal microscope (Carl Zeiss, LSM 780 under 10x magnification) and vertical serial images of tissue were captured at 5µm.

### 3.2 In *vivo* bioadhesive property of RPMNGel

To investigate the water resistance property of RPMN on dorsal mice skin, R6G@RPMNGel was applied topically on both diseased-free and psoriatic mice. After two minutes of incubation, the applied formulation was removed via gentle cleansing with a cotton swab, followed by continuous water irrigation to remove any remaining formulation from the skin and imaged using the IVIS Spectrum imaging system, Perkin Elmer, USA.

### 3.3. Assessment of permeation and retention behaviour of nanoparticles via IVIS system on diseased mice model

To understand nanoparticles’ permeation and retention behaviour in psoriatic and non-psoriatic mice, fluorescent dye rhodamine 6G (R6G) was used. Free R6G@Gel and R6G@RPMNGel were applied topically on dorsal mice skin. After 1 hour of application, gel was removed, cleaned, and imaged with an IVIS system. Again, mice were continuously treated for 24 hours; after 24-hour mice were cleaned and imaged, after which mice were not given any further treatment, and drug retention was checked for 48, 72, and 120 hours using 525/548 nm excitation/emission. After IVIS imaging, mice were sacrificed; their dorsal skin was collected, frozen in OCT solution, sectioned into 10 μm slices and imaged with a fluorescence microscope (LSM780, Carl Zeiss, Germany).

### 3.4. Establishment of imiquimod-induced psoriasis model

All experiments were carried out in compliance with the recommendation and permission of the Institutional Animal Ethics Committee (IAEC) (protocol number. SCI/IAEC/2022-2023/46), and studies were performed as per CPCSEA guidelines. Mice (8-12 weeks old) were acclimatized in typical laboratory conditions for one week before the start of the studies. Imiquimod (IMQ), a TLR ligand, has been applied topically and has been shown to cause skin inflammation equivalent to psoriasis. On experiment day, 62.5 mg of imiquimod cream (5%) was administered topically to the shaved region of the back for six days. The effectiveness of the treatment was evaluated by psoriasis area severity index (PASI), the most commonly used technique to assess the degree of severity based on thickness, erythema, and scaling [27]. On a scale of 0 to 4 i.e. 0, 1, 2, 3, and 4, which indicate nil, mild, moderate, severe, and very severe respectively. The cumulative scoring were also calculated, which includes erythema, scaling, and thickness, indicates the level of inflammation in a range of 0 to 12 [13].

### 3.5. *In vivo* therapeutic efficacy

Mice were randomly divided into six distinct groups (n = 6); group one was a negative control, whereas groups two to six received imiquimod topically to develop psoriasis-like lesions. In brief, group two served as a positive control, whereas groups three, four, five, and six administered free RAGel, BNLGel, RPMNGel, and commercialized clobetasol gel. All groups were treated consistently for six days, and effectiveness was evaluated daily utilizing the psoriatic area severity index (PASI) along with the thickness of the back skin by vernier calliper till day seven [28,29]. Finally, on day seven mice were euthanized, and their back skin, spleen, liver, heart, and kidney were surgically removed and preserved at −80°C for future experiments.

### 3.6 Spleen to body weight ratio

The spleen is a crucial immune organ involved in immunological reactions, phagocytosis, and eliminating old red blood cells, germs, or foreign antigens. An increase in spleen weight indicates stimulation proliferation and immune cell activation. At the end of the experiment, mice spleens were isolated, and their weight was recorded quickly to avoid errors in dehydration.

### 3.7 Drug safety assay

The *in vivo* cytotoxicity of the developed formulations in mice was evaluated by collecting primary tissues from different treatment groups, including back skin, spleen, liver, heart, and kidney, and observing daily body weight changes. Serum biochemical analysis was performed to determine the levels of Casein kinase (CKI), Alanine Amino Transferase (ALT), Aspartate Amino Transferase (AST), Blood Urea Nitrogen (BUN), and Creatinine in different groups.

### 3.8 Histopathological analysis

For histological study, tissues were dried, embedded in paraffin, stained with hematoxylin and eosin. The histological study examined various parameters like angiogenesis, supra papillary thinning, inflammatory infiltrates, munro’s microabscess, hyperkeratosis, parakeratosis, epidermal hyperplasia, inflammatory lesions, hyperkeratosis, parakeratosis, total skin damage score; liver parameters like inflammatory cell infiltration and hepatocyte degradation were studied; and spleen parameters like lymphocyte depopulation and splenocyte integrity of red and white pulp was studied.

## 4. Apparatus and chromatographic condition

HPLC samples were analyzed using a system equipped with an isocratic pump (Jasco 4180, Japan), an autosampler (Jasco 4050, Japan), and a UV detector (Jasco MD, 4015, Japan). A sensitive and reliable technique has been developed with an R^2^ of 0.99 throughout every experiment and a linear range of 2-60 µg/ml. The Agilent ZORBAX SB-C18 column (4.6 X 250 mm, 5um) was employed for separation. The mobile phase comprised 90:10 (v/v) methanol and water, with an injection volume of 20 μl and a 1.0 mL/min flow rate. The UV detection wavelength was 278 nm. Data was acquired using the Chrom Nav chromatographic data system program.

## 5. Statistical Analysis

Three independent experiments were given as mean ± standard deviation or standard error of the mean (SEM). GraphPad Prism software (version 8, San Diego, California, USA) analyzed group differences using ANOVA or student t-tests. A p-value of <0.05 was considered statistically significant. Asterisks indicate p-values in figures (**p <0.01,***p <0.001,****p <0.0001).

## 6. Result and discussions

### 6.1 Rapamycin NP (RPMN) loaded gels (RPMNGel) have better material characteristics, controlled drug release and long-term stability

Hydrodynamic particle size, surface charge, polydispersity index, release profile and encapsulation efficiency were significant factors in determining the most suitable formulation. The particle size of optimised RPMN was approximately ≈277.6 nm **(Fig. 1A)** with a spherical shape **(Fig.1B)** and PDI of 0.21 indicates a uniform particle size distribution. The zeta potential **(Fig.1C)** was ≈ −17.5 Mv, showing good colloidal stability and the amount of drug encapsulated inside nanoparticles was ≈83%. RPMNGel was fabricated by combining RPMN with carbomer overnight and stirring to form RPMNGel. Rheological data **(Fig 1.D)** of RPMNGel revealed the viscosity of 22079.8 cp at 37°C and viscosity vs shear rate and shear stress vs shear rate displayed that the gel showed non-newtonian behaviour, demonstrated by its thixotropic behaviour and varied shear thinning properties. The texture analyser data revealed that the gel has high adhesiveness and cohesiveness, which is better for skin adherence **(Fig. 1E, Table Supp.2).**

**Fig. 1:**
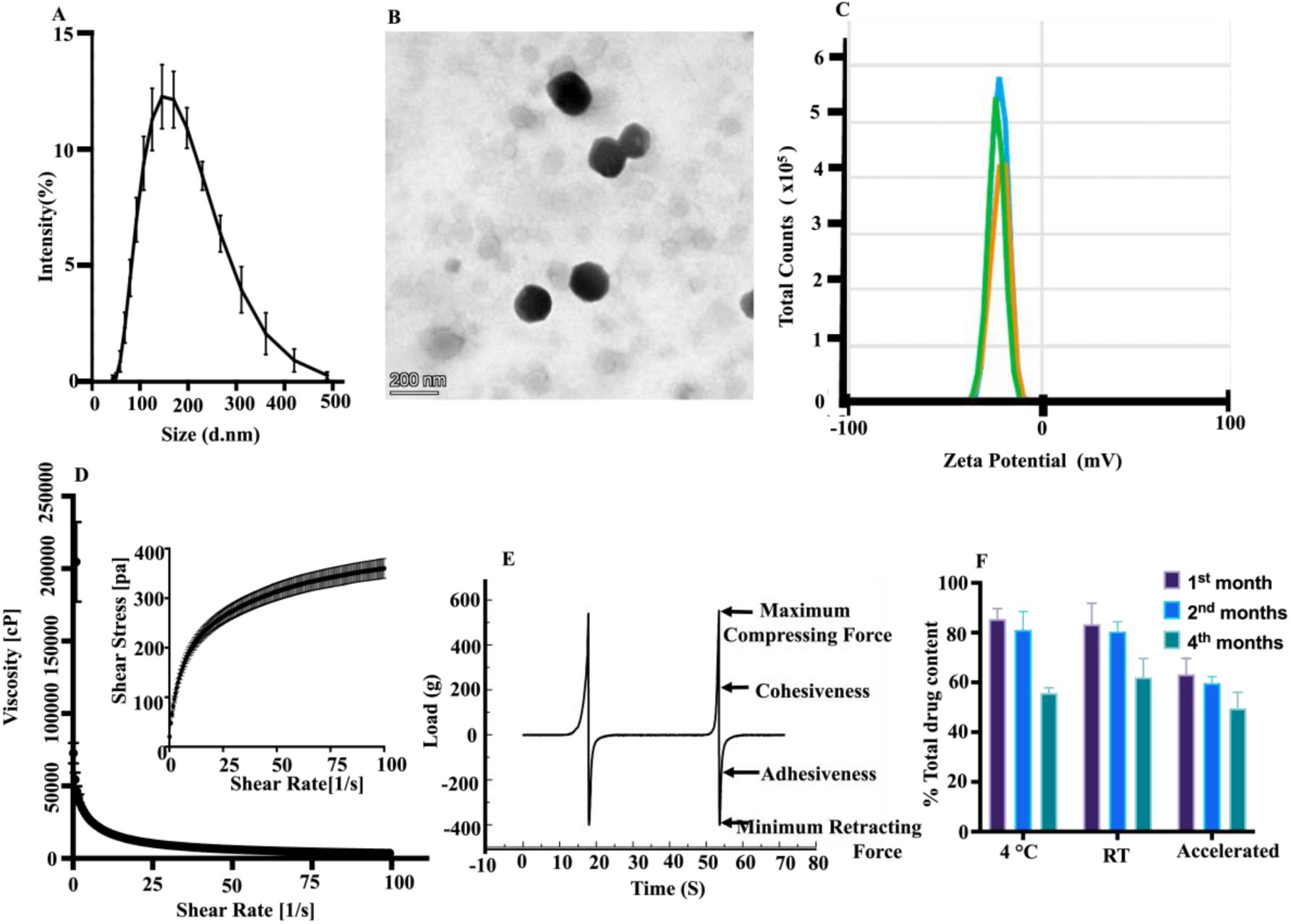
Physiochemical Characterisation of RPMN & RPMNGel. (A)Average hydrodynamic particle size distribution. (B)TEM image. (C)Zeta potential. (D) shear stress vs shear rate and viscosity versus shear rate graphs of RPMNGel. (E) Textural profile of the RPMNGel. (F) Stability study under various storage conditions. Data are represented as mean ± SD (n = 3)

In vitro release profiles of RPMNGel have been shown therein **(Fig. Supp 1).** The data indicated that release of rapamycin from RPMNGel showed prominent sustained release for 48 hours with no burst release. This extended-release might be adhering RNPL to or being embedded in the tiny pores of carbomer gel, making diffusion difficult and preventing burst release. Antioxidant properties were also investigated via DPPH assay, which showed that RPMN has free radical scavenging activity of ≈55.9% after 10 minutes of incubation in DPPH solution, proving better free radical scavenging ability.

The stability study of RPMNGel showed similar drug content at 4°C and RT but decreased in the 4th month, possibly due to drug degradation. Concentration of drug at accelerated storage condition significantly reduced in the first month, possibly due to higher temperatures and humidity. No changes were observed in terms of syneresis, consistency, colour change, phase separation, or grittiness during the entire duration of the study **(Fig 1.F)**.

### 6.2 RPMNGel can easily penetrate inside psoriatic skin

Franz diffusion apparatus was used to study *in vitro* transdermal skin permeation and deposition of RPMNGel and free RAGel on excised psoriatic mice skin (**Fig. 2A)**. The results suggested a more substantial accumulation of rapamycin in both the dermis and epidermis from nanoparticle gel nearly two times higher than the free rapamycin gels (**Fig. 2B&C)**. This could be due to the occlusive effect of RPMNGel, which forms an intact thin coating on the skin’s surface, or the accumulation of skin appendages such as hair follicles, which may serve as a long-term drug repository. However, a negligible amount of drug was penetrated inside the donor compartment.

**Fig. 2:**
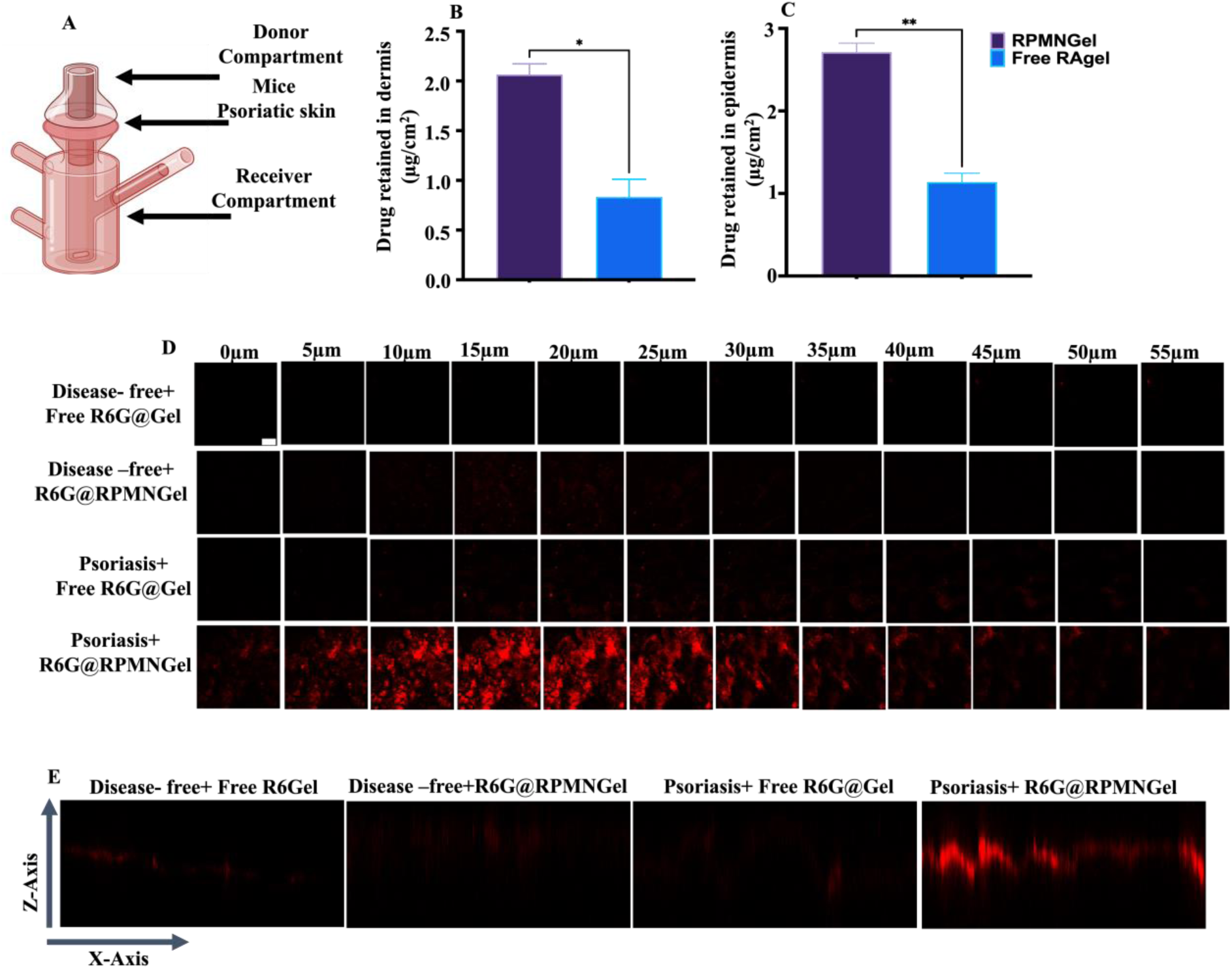
*Ex vivo and in vivo* penetration and deposition of drug and R6G dye.(A) experimental setup with the Franz diffusion apparatus.(B&C) amount of drug detected in the epidermis and dermis.(D) representative confocal serial microscopic imaging of mice ear skin treated with R6G@Gel or R6G@RPMNGel on disease-free and psoriatic mice, images were obtained at 5 µm intervals on skin surfaces (Scale bar, 153 µm). (E) X-Z orthogonal image of psoriatic and disease-free mice treated R6G@ Free gel and R6G@BNPGel. Data are represented as mean ± SD (n = 3). P values were determined using a t-test. **p <0. 01,***p <0. 001.

Similarly, in support of these observations, RPMNGel was labelled with the fluorescent dye rhodamine 6G, allowing tracking and exploring the permeation process inside the psoriatic skin by comparing with control i.e. free rhodamine 6G hydrogel. After 24 hours of application, ear samples were imaged with a CLSM and it was observed that in healthy ear skin, neither Free R6G@Gel nor R6G@RPMNGel displayed any fluorescence signal, as shown in (**Fig. 2D)** and their quantitative distribution **(Supp.Fig. D)**. However, R6G@RPMNGel demonstrated significant fluorescence in psoriatic skin, reaching depths of up to 45μm [31,32]. This depth allowed RPMN to pass through the psoriatic skin layers and reach the epidermis, which could be acquired by keratocytes or infiltrating lymphatic cells to function. The x-z axis orthogonal view of the skin at each interval of 5 μm was depicted (**Fig. 2E),** and the outcomes provided additional evidence of the increased penetration of R6G@RPMNGel in psoriatic skin along with the uniform distribution throughout the skin. The result concluded that R6G@RPMNGel could easily penetrate the deeper layer of skin and impart therapeutic activity.

### 6.3. RNPL ameliorates psoriatic symptoms in an IMQ-induced mouse psoriasis model

We subsequently evaluated the therapeutic efficacy of prepared nanoparticles on IMQ-induced psoriatic model in mice’s backs. These models have been recognised as accurate preclinical psoriasis models as it exhibit inflammation closely related to human psoriasis, such as a thickened epidermis, abnormal keratinocyte-related protein expression, inflammatory cell infiltration, and the secretion of proinflammatory cytokines [33,34]. **(Fig. 3A&B)** represent the timeline and duration of application of IMQ with different formulations with a gap of 8 hours and a photographic image of mice on the 1^st^ to last days. The severity and therapeutic efficacy were analysed based on the PASI score. The topical administration of IMQ on the back skin for six consecutive days developed psoriatic-like lesions such as skin thickening, scaling, and erythema, PASI score respectively **(Fig. 3C)**. In the positive control, significant psoriatic symptoms like scaling, erythema, and thickening started to increase progressively from day 1 to 7. Their cumulative score reached 10.25 on day 7; similarly, in the treatment of RPMNGel, the disease severity was lessened, and the PASI value reduced to 1.75, which showed that RPMNGel showed a better therapeutic role in ameliorating psoriasis as compared to that of other treatment group.

**Fig. 3:**
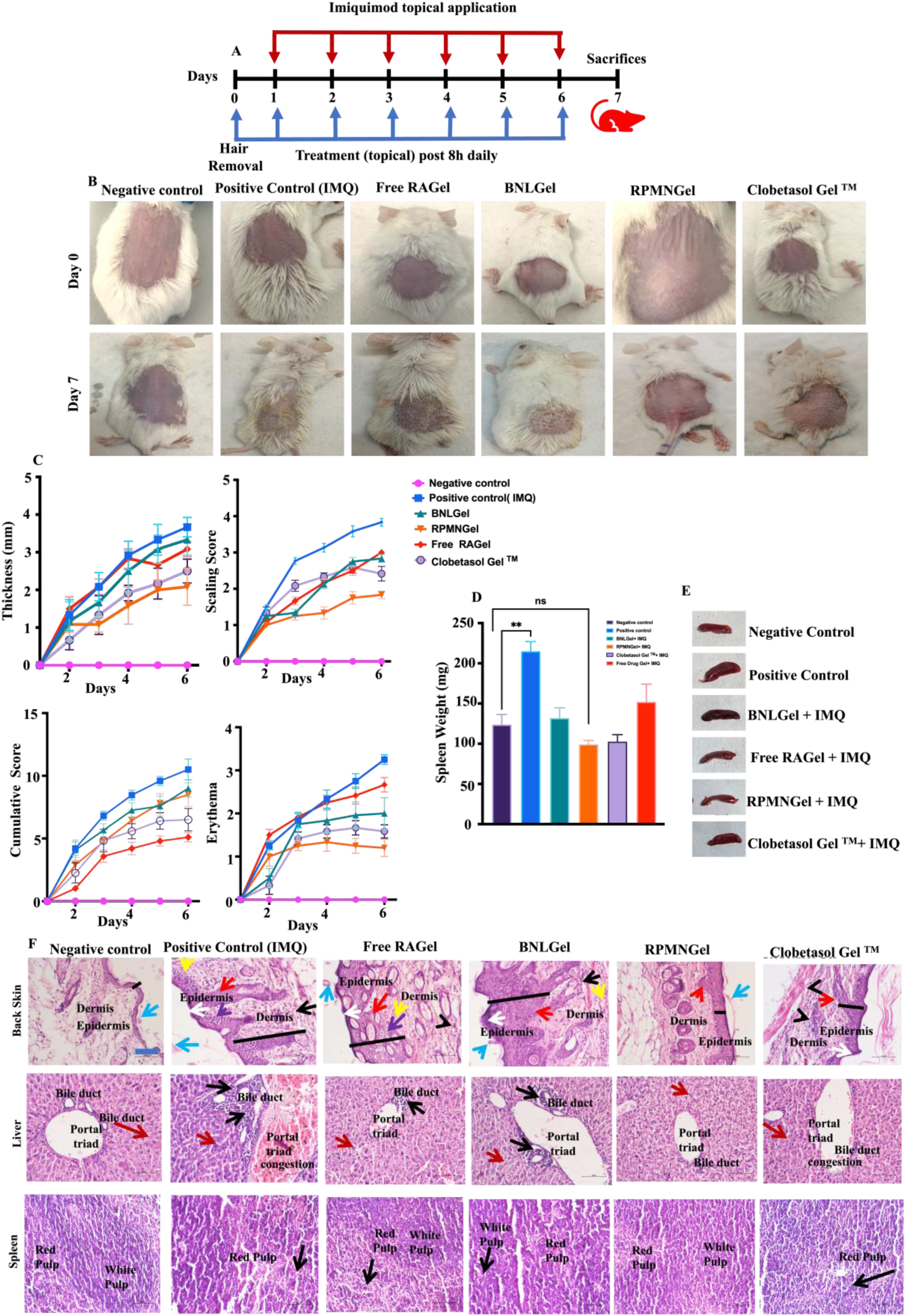
RPMNGel ameliorates imiquimod-induced psoriatic-like skin lesions in mice models. (A&B) Phenotypical representation of mice’s back skin and their IMQ application and treatment duration. (C)Thickness, scaling, erythema were calculated based on the clinical PASI score on a scale of 0–4, and the total cumulative score between (0–12). (D&E) Images of the spleen and the average weight of various treatment groups. (F) Histological evaluation of back skin, liver and spleen (Scale bar, 10µm) **Liver**: *Black Arrow*-Inflammatory cells, *Red Arrow*-Hepatocyte **Spleen***: Black Arrow*-Degenerate Lymphocyte **Skin**: Keys: *Blue arrow*-Parakeratosis, *Red Arrow*-Epidermal hyperplasia, *White Arrow*-Hyperkeratosis, *Black arrow*-Inflammatory cells, *Black Line*-epidermal thickness, *yellow arrow*-Haemorrage, *Arrow head*-Neo-angiogenesis*, Violet arrow*-Epidermal edema (microabcess)

The application of IMQ substantially increased the size and weight of the spleen from ≈123.7 mg to ≈215.0 mg (**Fig. 3D&E)**, mainly due to the inflammation and accumulation of cells in the spleen, increasing the spleen’s size and weight. The spleen weight was significantly reduced in the group treated with RPMNGel, which was almost similar to the negative control.

To further prove the hypothesis, histological study of the back skin, spleen, and liver were studied **(Fig. 3F)** and the total skin damage was noted. When comparing RPMNGel, less skin damage was observed as compared to other treatment groups (**Table 1**), indicating the efficacy of RPMNGel to reduce the expression of various psoriatic factors.

**Table 1.** indicates multiple signs reflecting the degree of skin injury following topical application of different treatment groups on the Back skin imiquimod-induced psoriatic model. Scoring (+ Indicate Mild), (++ Indicate Moderate), (+++ Indicate Severe)

| Group | Parakeratosis | Hyperkeratosis | Inflammatory cells | Epidermal Hyperplasia | Capillary proliferation (Neo-angiogenesis) | supra papillary thinning | Micro-abscess (Odema) | Pustule of Kogoj | Total Skin Damage Score |
| --- | --- | --- | --- | --- | --- | --- | --- | --- | --- |
| Negative Control | 0 | 0 | 0 | 0 | 0 | 0 | 0 | 0 | 00 |
| Positive Control | + | ++ | +++ | +++ | + | +++ | ++ | + | 16 |
| BNLGel | + | ++ | ++ | +++ | + | +++ | + | 0 | 13 |
| RPMNGel | + | 0 | 0 | + | + | + | 0 | 0 | 04 |
| Free RAGel | + | ++ | + | + | ++ | + | +++ | 0 | 11 |
| Clobetasol<sup>TM</sup> Gel | + | 0 | ++ | ++ | + | + | 0 | 0 | 07 |

Similarly, histology of the liver revealed normal portal triad, central vein, hepatocyte having round to oval nucleus in negative control, RPMNGel treated group and clobetasol Gel^TM^. However, in positive control, BNLGel and free RAGel showed inflammatory cells infiltration in peripheral of bile duct. Interestingly, Histology of spleen showed normal architecture of white pulp and red pulp of tissue in negative control and RPMNGel group and remaining other group showed degenerative lymphocyte in red pulp. These results further confirmed that RPMNGel caused less liver, spleen and skin damage than other groups.

### 6.4 RPMNGel showed bio adhesive and better skin deposition properties

To assess the skin adhesive property of RPMN, the R6G@RPMNGel was applied topically on the dorsal side of healthy and diseased mice and allowed to incubate for 2 minutes. After repeated water irrigation, followed by gentle wiping, imaged with IVIS which showed the presence of fluorescence signal in diseased mice as compared to healthy mice (**Fig. 4A, Supp Fig. B)**. This might be attributed to R6G@RPMNGel not adhere on healthy skin and the restricted binding to the most superficial components of intact normal stratum corneum [35].

**Fig. 4:**
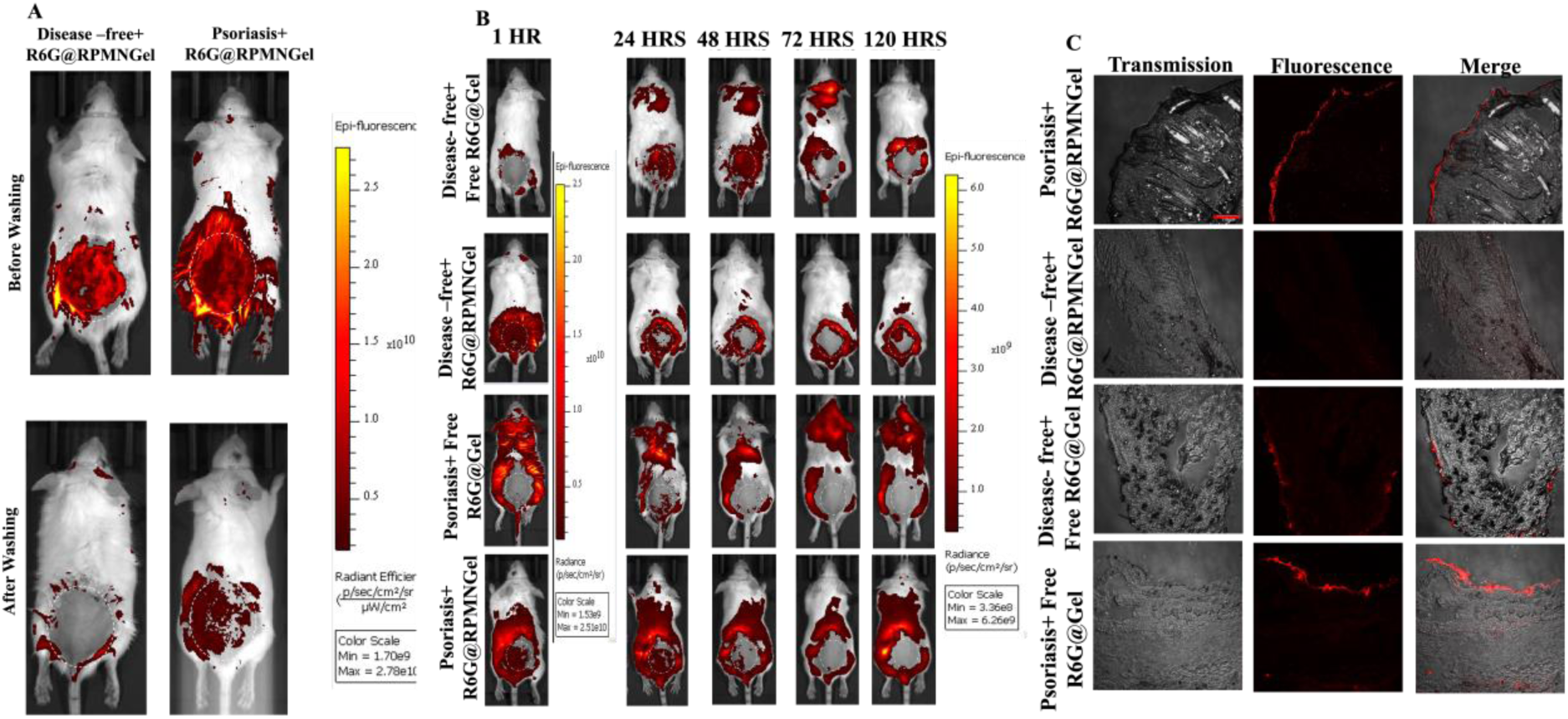
Bioadhesive, penetration and excretion profile of RPMNGel. (A) Comparison of skin adhesive property of R6G@RPMNGel. (B&C) Permeation, excretion and their retention profile of free R6G@Gel or R6G@RPMNGel in the back skin tissue. (Scale bar, 110 µm)

However, psoriatic skin has a damaged skin layer, therefore R6G@RPMNGel may adhere to the skin properly and show better penetration profile. This showed that RPMN could be used as promising tool for treatment of psoriatic skin due to its better penetration profile to the skin.

The relative skin deposition and excretion profile of free R6G@Gel and R6G@RPMNGel in both diseased and non-diseased groups was also analysed. It was observed that 24 hours post-treatment, the fluorescence intensity was higher in the diseased group treated with R6G@RPMNGel than in other groups. By day four, less fluorescence signal was observed this might be due to the entirely removal of nanoparticles from the body (area depicted in the white circles) (**Fig. 4B Supp Fig. C)**. This decrement was observed more in the healthy control group than diseased nanoparticles treated model due to the incapability of the nanoparticles to adhere to the epidermis or its non-bioadhesiveness or might be partially attributed to the skin renewal. On a similar note, to estimate dye retention on the skin, the skin was collected, sliced and imaged with a fluorescence microscope. It was concluded that psoriatic skin exhibited a higher accumulation of R6G@RPMNGel than the healthy group (**Fig. 4C Supp Fig. D)**. However, in the healthy group, formulations were attached to the skin surface without penetrating the epidermal layer. According to these findings, R6G@RPMNGel may penetrate psoriatic-affected regions and is primarily retained for extended periods, which could mitigate adverse effects on healthy skin.

### 6.5 Topical application RPMNGel is safe confirmed by Histological study and biochemical assay

Developing a new treatment regime to treat any disorders requires prioritizing safety concerns. To assess possible toxicity, various experiments, such as *in vitro* and *in vivo* cytotoxicity, were carried out in human keratinocyte cell lines and swiss albino mice for 24 and 72 hours, respectively. *In vivo* toxicity of RPMNGel, BNLGel, and free RAgel was studied using H&E staining, while PBS was kept as a control. All the treated groups did not showed any sign of toxicity as compared to the control group **(Fig. 5A**). Similarly, *invitro* study on HaCaT cells showed that RPMN inhibited proliferating cells more effectively than BNL and free RA in DMSO **(Fig. Supp E)**.

**Fig.5:**
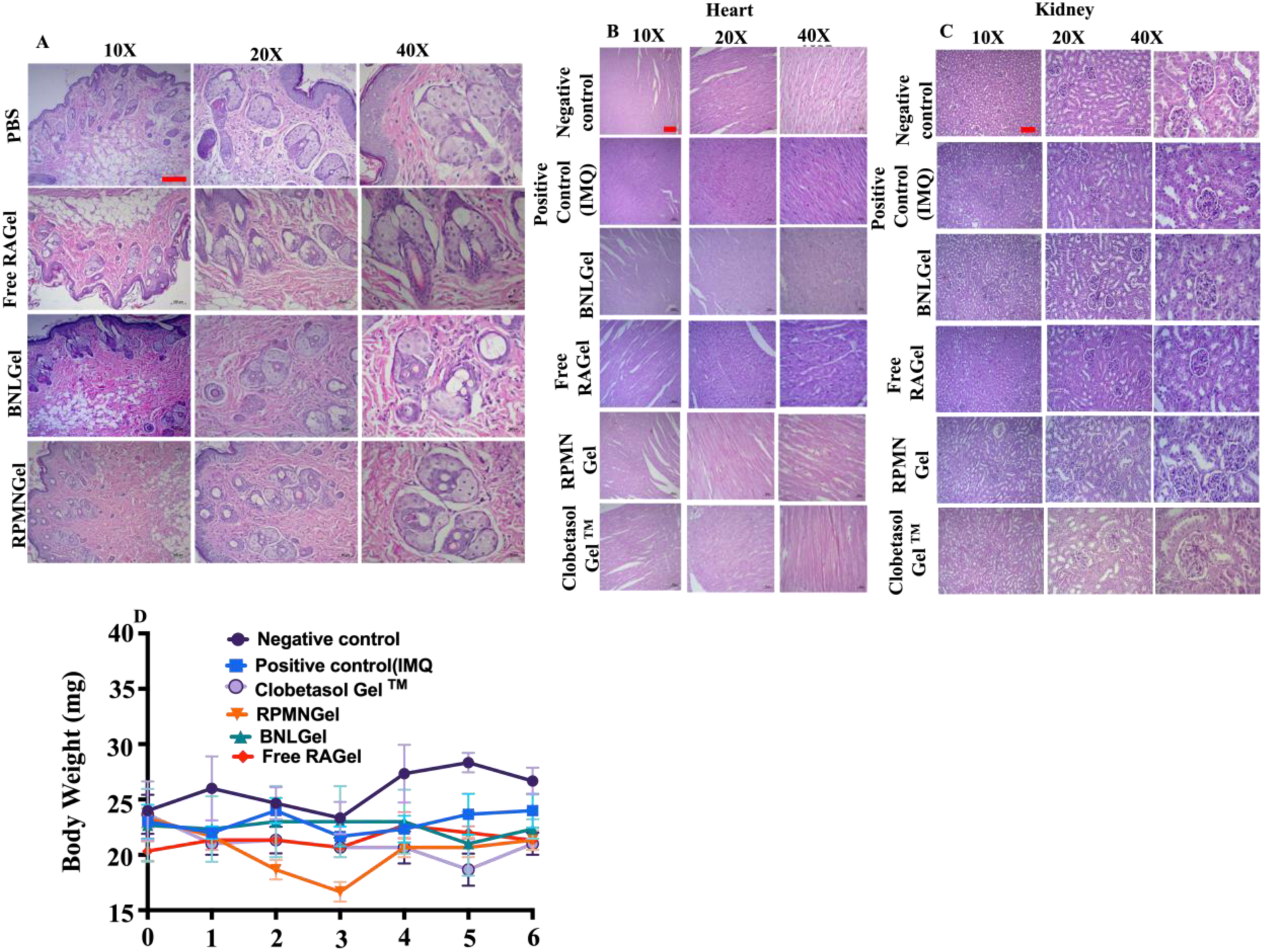
Drug safety assay. (Fig. A) indicates *in vivo* cytotoxicity on Swiss albino mice, (B&C) H&E of heart and kidney. (D) Body weight % of treatment groups (Scale bar 10X-100 µm, 20X-50 µm, 40X-20 µm).

Similarly, at the end of the study, the heart **(Fig. 5B**) and kidney **(Fig. 5C**) were collected for H&E staining to understand any adverse effects on the organ and no major or minor signs of significant damage were observed compared to the positive control, which enhanced the particle safety for application.

In **(Fig. Supp F)** serum biochemical studies were conducted to understand the significant changes in critical organs, including the liver and kidney. All the groups showed notable elevations in the liver enzyme levels, like aspartate aminotransferase (AST), alanine aminotransferase (ALT) levels, casein kinase I, however, groups treated with RPMNGel and clobetasol gel^TM^ gel showed little changes in these enzymes as compared to a negative control. Meanwhile, BUN (blood urea nitrogen) did not showed difference in all the groups. Level of CRE (creatinine) was similar as compared to negative control and elevated in the diseased group and other group.

Additionally, the average body weight of the mice has been monitored since the first day. However, the body weight of the negative control and other treatment groups did not exhibit significant changes compared to the negative control **(Fig. 5D**).

## 6. Conclusions

In the present work, we confirmed that PI3K/Akt/mTOR inhibitors can be used to control and manage psoriasis. RPMN can easily permeate through the skin barrier and retained in the various skin layers with negligible systemic absorption. *In vivo* study concluded that the nanoparticle has skin adhesive properties and can only target psoriatic skin but not healthy skin, along with the required therapeutic efficacy for psoriasis treatment. Drug safety assays like biochemical assay histological staining of significant organs suggested that the formulation is safe without side effects. Thus, rapamycin-loaded lipid polymer conjugated nanoparticles can be used as an alternative therapy for psoriasis treatment with better skin permeation and adhesiveness. Though the information collected is critical in assessing its effectiveness, more clinical trials and very long-term stability studies are also necessary to support the outcomes.

## 7. Authors Contributions

**RK-** Conceptualization, methodology, execution, investigation, analyzing experiments, literature survey, writing original draft, review & editing, **RB –** particle characterization and optimization, review and editing, **SR -** Histology slide preparation and interpretation and IVIS imaging., **Late RB-** Initial fund acquisition, supervision and conceptualization, **SS&RS**-Conceptualization, Supervision, Investigation, Project administration, Funding acquisition, Resources, review & editing of manuscript

## 8. Acknowledgement

The author acknowledges SAIF (Sophisticated Analytical Instrument Facility) and IRCC (Industrial Research & Consultancy Centre) at the Indian Institute of Technology Bombay, India, for providing instrumental support. Roshan Keshari also acknowledges the DAAD (German Student Exchange Programme), Govt. of Germany, for a research fellowship.

## 9. Declaration of competing interest

All authors declare no conflicts of interest.

## Notes

### Competing Interest Statement

The authors have declared no competing interest.

### Summary of Updates

There were some issues with the images, which was resolved.

