## Supplementarty for "Transcutaneous delivery of disease-specific PI3K/Akt/mTOR inhibitor-based hybrid nanoparticles in hydrogel system for the management of psoriasis: Insights from *in vivo* studies"

**^1#^ Deceased on 8^th^ July 2021**

***Corresponding authors**

**Department of Biosciences and Bioengineering,**

**Indian Institute of Technology (IIT) Bombay,**

**Powai, Mumbai, India,**

**, ***

**Table T1 -Test method used for textural analysis of EUNPGel**

| **Parameters** | **Value** |
| --- | --- |
| Test Type: | TPA |
| Target: | 36.0 mm |
| Hold Time: | 0 s |
| Trigger Load: | 0 g |
| Test Speed: | 2.00 |
| Return Speed: | 2 mm/s |
| # of Cycles | 2.0 mm/s |
| Target Type | Distance |
| Recovery Time: | 0 |
| Same Trigger: | True |
| Pretest Speed: | 2.00 mm/s |
| Data Rate: | 30.00 points/sec |
| Probe: | TA2/1000 |
| Fixture: | TA-BT-KIT |
| Load Cell | 10000g |

**Table 2. Texture Analysis profile RPMNGel Data are represented as mean ± SD (n = 3)**

| **Hardness (g)** | 521.2±20.7 |
| --- | --- |
| **Adhesive force (g)** | 390.3±11.2 |
| **Adhesiveness (mJ)** | 8.1±0.7 |

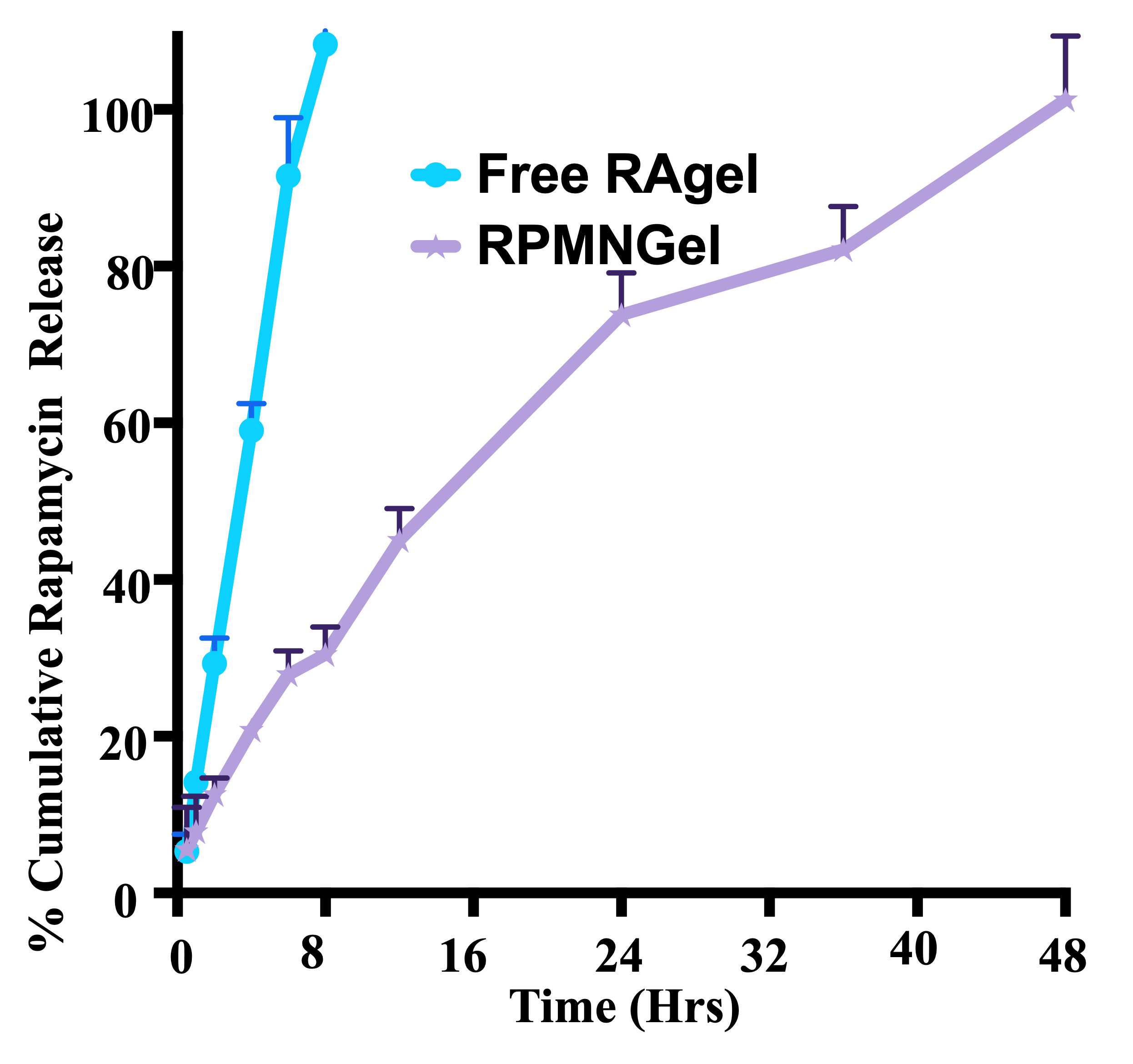

(A) ***In vitro* cumulative release profile of eugenol from RPMNGel and Free RAgel**

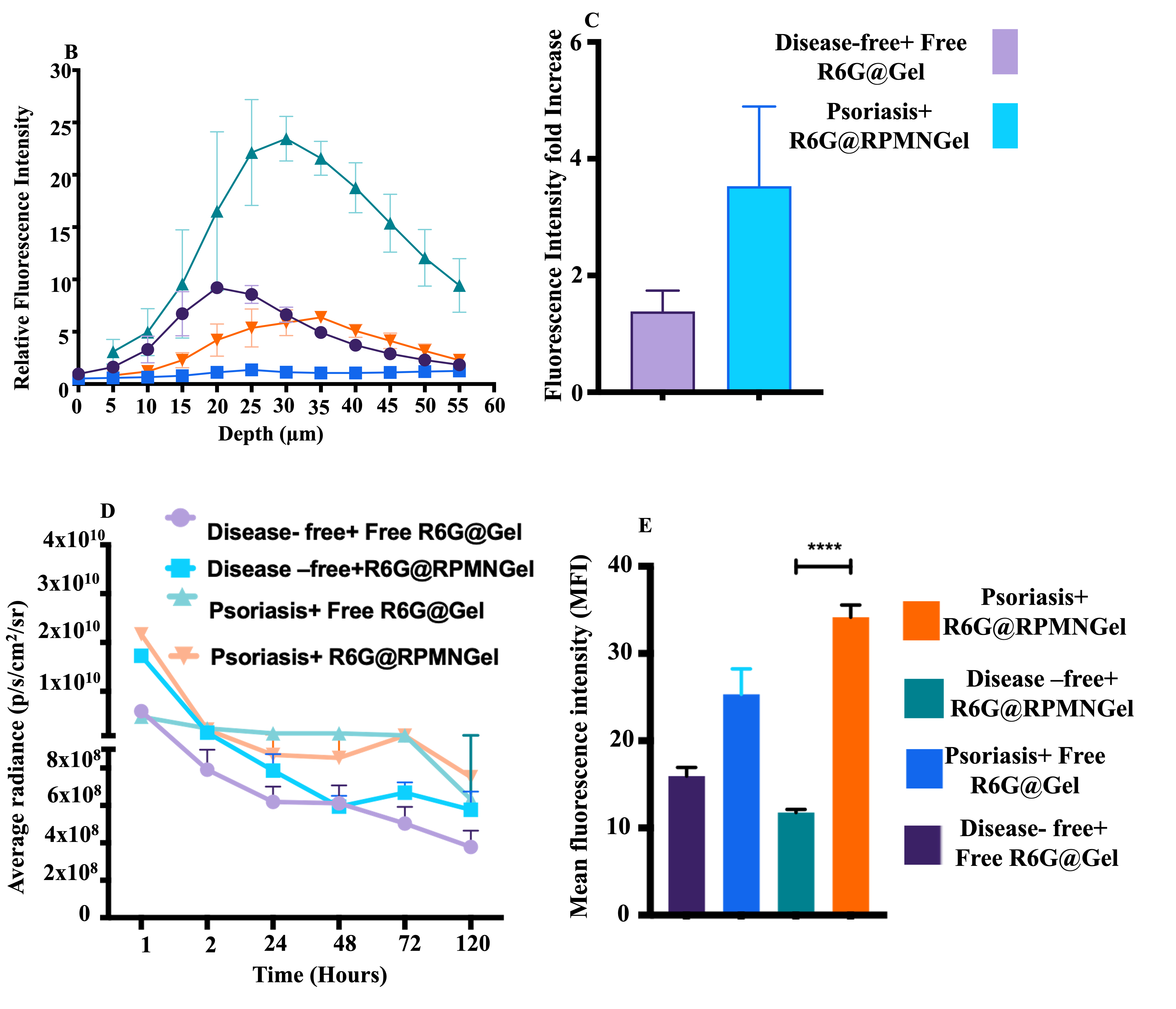

**Fig. (B,C,D&E) Quantification of Bioadhsive, penetration and excretion profile of RPMNGel of on both diseased and non-diseased group treated with free R6G@Gel and R6G@RPMNGel. (B) permeation of R6G labelled RPMNGel. (C) Quantification in fluorescence intensity fold changes in bio adhesive profile of RPMNGel. (D) Average radiance in skin penetration and excretion profile. (E) quantification of dye retention and their quantification. Data are represented as mean ± SD (n = 3). P values were determined using a t-test. **p <0. 01,***p <0. 001.**

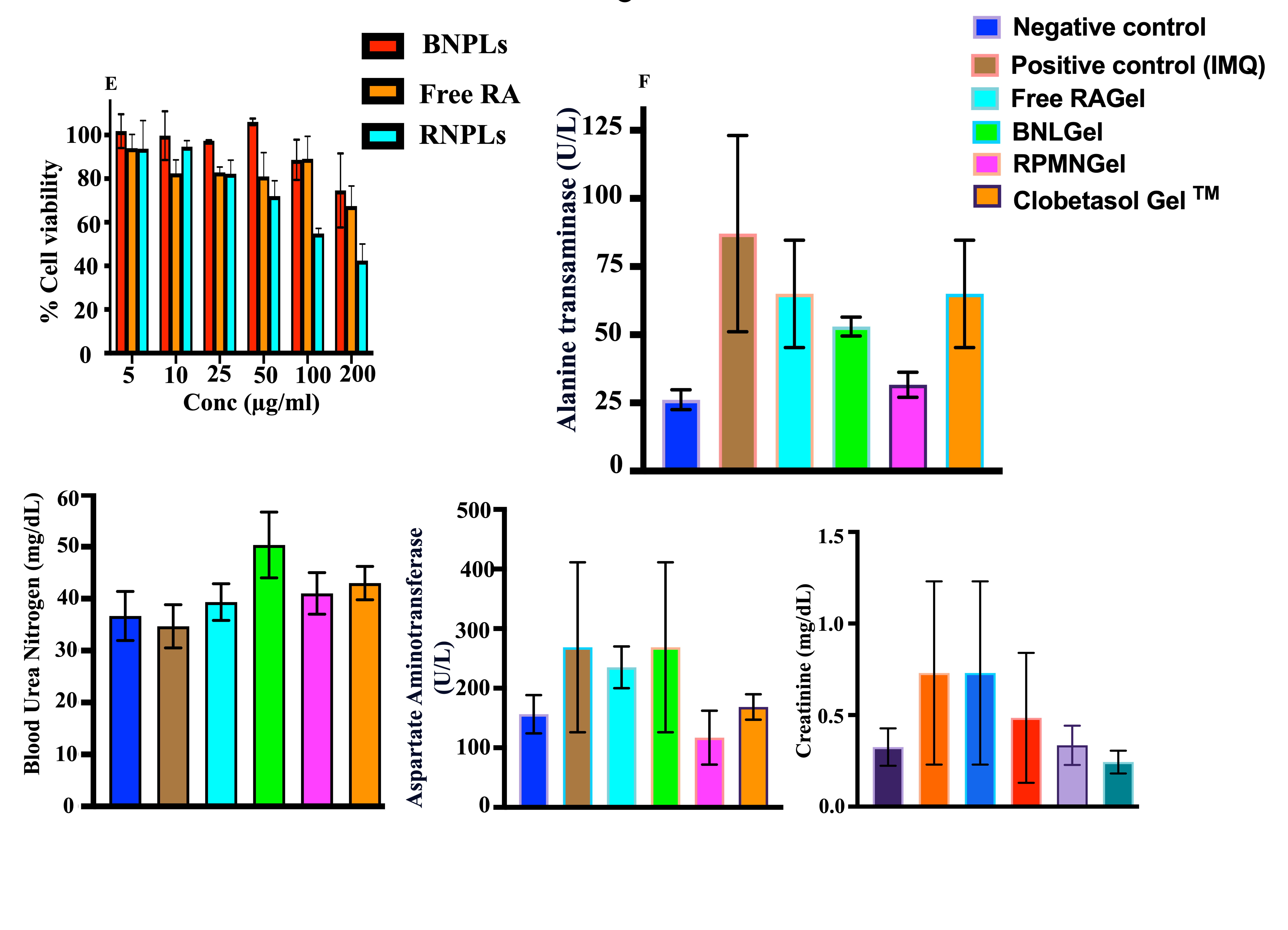

**(Fig. E&F) Evaluation of and *Ex vivo* ,*In vivo* cytotoxicity. (E) Cytotoxicity on HaCat cell by MTT assay. (F) Biochemical analysis of Alanine Amino Transferase (ALT), Blood Urea Nitrogen (BUN), Aspartate Amino Transferase (AST), Creatinine were measured in all treatment groups (Magnification: 20X). Data are represented as mean ± SD (n = 3). P values were determined using a t-test. ****p <0. 0001, ***p <0. 001, **p <0. 01.**
